# A Nile Grass Rat Transcriptomic Landscape Across 22 Organs By Ultra-deep Sequencing and Comparative RNA-seq pipeline (CRSP)

**DOI:** 10.1101/2022.02.04.479193

**Authors:** Huishi Toh, Atefeh Bagheri, Colin Dewey, Ron Stewart, Lili Yan, Dennis Clegg, James A. Thomson, Peng Jiang

## Abstract

RNA sequencing (RNA-seq) has been a widely used high-throughput method to characterize transcriptomic dynamics spatiotemporally. However, typical RNA-seq data analysis pipelines depend on either a sequenced genome or corresponding reference transcripts or them both. This restriction makes the use of RNA-seq for species lacking both of sequenced genomes and corresponding reference transcripts challenging. Nile grass rat (*Arvicanthis niloticus*) is a diurnal rodent species with several unique characteristics making it as a useful model to study diet-induced type 2 diabetes and other physiological or behavioral processes due to its diurnal nature. However, there is neither a high-quality annotated Nile grass rat genome nor a reference transcript sets available so far, making it technically challenging to perform large-scale RNA-seq based transcriptomic studies. Although we are working on the first draft of Nile grass rat genome, a well annotated genome typically requires several rounds of manually reviewing curated transcripts and can take years to achieve. To solve this problem, we developed a Comparative RNA-Seq Pipeline (CRSP), integrating a comparative species strategy but not depending on a specific sequenced genome or species-matched reference transcripts. Benchmarking suggests the CRSP tool can achieve high accuracy to quantify gene expression levels. In this study, we generated the first ultra-deep (2.3 billion × 2 paired-end) Nile grass rat RNA-seq data from 59 biopsy samples representing 22 major organs, providing a unique resource and spatial gene expression reference for using Nile grass rat as a model to study human diseases. To facilitate a general use of CRSP, we also characterized the number of RNA-seq reads required for accurate estimation via simulation studies. CRSP and documents are available at: https://github.com/pjiang1105/CRSP.

**Highlights:**

- CRSP is a novel software tool which can quantify gene expression levels from RNA-seq data for species lacking both a sequenced genome and corresponding reference transcripts.
- Nile grass rat is a unique diurnal rodent species (day active but not night active) with several unique characteristics making it as a useful model to study diet-induced type 2 diabetes and other physiological or behavioral processes due to its diurnal nature.
- We generated the first ultra-deep (2.3 billion × 2 paired-end reads) Nile grass rat RNA-seq data from 59 biopsy samples representing 22 major organs, providing a unique resource and spatial transcriptomic reference (e.g., tissue gene expression baseline) for using Nile grass rat as a model to study human diseases.

## 1. Introduction

RNA sequencing (RNA-seq) has been widely used for characterization of transcriptomic dynamics[1] with different biological contexts, such as development and diseases. Current widely used RNA-seq tools either rely on an annotated genome or species matched reference transcripts (e.g., RSEM [2]). For species lacking both of sequenced genomes and corresponding reference transcripts, performing RNA-seq data analysis remains challenging. In general, there are two alternative computational strategies which can potentially solve this problem. The first strategy is to estimate gene expression values via directly mapping RNA-seq reads to a well-annotated genome of a closely related species [3]. This strategy works well if the genomic DNAs of two species are very similar. However, it is unknown that for two species, how similar is similar enough to allow such cross-species RNA-seq reads mapping. Given that not all species can find a closely related species with a well-annotated genome, this strategy is only limited to a few cases. The second strategy, instead of applying a direct cross-species RNA-seq reads mapping, is to assembly RNA-seq short reads to a longer contigs set first and then annotate these contigs via mapping them to a protein database of another species [4]. The gene expression values were further imputed via re-mapping RNA-seq reads to contigs and aggregating contigs level expression to a gene level expression [4]. The basic assumption of this strategy is that the majority of genomic mutations in two species are non-coding or synonymous (do not change amino acids). Hence combing *de novo* transcripts assembly [5] with cross-species comparative mapping contigs to a protein database can partially solve the problem regarding to using a distantly but not necessarily closely related species as a reference [4]. Our prior studies suggest this strategy works well even for two species with divergence time ∼ 300 MYA (million years ago), such as estimating axolotl gene expressions (RNA-seq data) via using a frog protein database [4]. Although this strategy looks promising, implementing such strategy in a particular project is very complicated [4]. There is lacking of software tools which can implement this strategy into a user-friendly generalized pipeline. Furthermore, the only evidence to show this strategy works well was based on comparing multiple key marker gene expression patterns with prior knowledge [4]. Since there is no benchmarking of these approaches, and thus it is largely unknown that how well this strategy actually performs.

To facilitate a general use of RNA-seq data to species without any prior annotated references, we developed CRSP (Comparative RNA-seq pipeline) tool which is conceptually similar the second strategy. CRSP integrates a set of computational steps, such as mapping *de novo* transcriptomic assembly contigs to a protein database, using Expectation-Maximization (EM) algorithm to assign reads mapping uncertainty, and integrative statistics to quantify gene expression values. Our evaluations of CRSP on human RNA-seq data but use mouse proteins as a reference show that its estimated gene expression values are highly correlated with gene expression values estimated by directly mapping to the human genome. We also estimated the number of RNA-seq reads required for accurate estimation via simulation studies. Our simulations suggest that 10 to 20 million single-end reads are sufficient to achieve reasonable gene expression quantification accuracy while a pre-compiled *de novo* transcripts assembly from deep sequencing (although not required) can dramatically decrease the minimal reads requirement for the rest of RNA-seq experiments.

Nile grass rat (Arvicanthis niloticus), native to East Africa, is one of a few diurnal (day active) rodent models established for laboratory study. Unlike commonly used laboratory rodents, such as mice, rats and hamsters, which are nocturnal (active at night), Nile grass rats are diurnal as humans. There are fundamental differences between diurnal and nocturnal species in the temporal organization ranging from behavior, physiology to neural activity and gene expression, which are all under the control of the circadian time-keeping system. The organization of circadian systems in nocturnal versus diurnal species differs in a more complex manner than a simple flip in the daily pattern [6]. Therefore, a diurnal model has unique translational value in biomedical research and will contribute to a better understanding of mechanisms underlying health and disease in diurnal humans. This diurnal rodent has a natural diet primarily of leaves and stems, supplemented with insects, seeds, and fruits. However, when fed a diet with a higher caloric content, it consistently develops type 2 diabetes (T2D). Unlike other laboratory rodents, which require high-fat feeding and take a long time to develop T2D, Nile grass rat develop T2D rapidly (8-10 weeks)[7] with high-carbohydrate diets. These features make Nile rat as a prominent T2D model. However, transcriptomic studies in Nile grass rat are very limited because its genome is neither sequenced nor annotated [8] making RNA-seq data analysis challenging. Although we are working on the first draft of Nile grass rat genome sequencing, it may still take several years from the first sequenced genome to a high quality and well annotated genome which can be used as a reference for a typical RNA-seq data analysis [9, 10]. To further demonstrate CRSP is a useful tool which can study transcriptome without relying a corresponding sequenced genome or corresponding reference transcripts, we generated an ultra-deep (2.3 billion × 2 paired-end) Nile grass rat RNA-seq data from 59 biopsy samples representing 22 major organs. Using CRSP, we generated the first Nile grass rat gene expression BodyMap, providing a unique resource and spatial transcriptomic reference (e.g., tissue gene expression baseline) for using Nile grass rat as a model to study human diseases.

## 2. Materials and methods

### 2.1 Comparative RNA-seq Pipeline (CRSP)

CRSP takes RNA-seq raw reads and the transcriptomic assembly contigs (e.g., generated by Trinity [5]) as input, and then integrate a set of computational steps to estimate gene expression levels. The CRSP workflow contains the following steps:

a. Merging assembly (Contigs): the contigs from different tissues, conditions or batches are merged and then reduced to a non-redundant contig set;
b. Mapping contigs to a protein database: the non-redundant transcriptomic contig set is mapped to a protein database (e.g., human or mouse protein database) via the NCBI BLASTX tool [11]. CRSP excludes hits with E-value >= 10^−5^, and the best hit is selected for each contig;
c. Mapping RNA-seq reads to contigs: The RNA-seq reads are mapped to contigs, and then a contig level abundance estimation is performed via RSEM [2];
d. For each RNA-seq sample, gene-level RNA-seq mapped read counts are obtained by summing the counts of the contigs that are mapped to proteins with the same gene symbol; (e) Relative abundances, in terms of transcripts per million (TPM), for genes are computed by first normalizing each gene’s fragment count by the sum of the “effective lengths” (length less the read length) of the contigs mapped to that gene and then scaling the resulting values such that they sum to one million over all genes.

A more detailed workflow of CRSP can be found in **Supplementary Figure S1**.

### 2.2 RNA-seq datasets used for benchmarking

#### (a) Human single-end RNA-seq dataset

The single-end reads (human, embryonic stem cell (ES), H1 cell line) was from our published work [12].

#### (b) Human paired-end RNA-seq dataset

We also generated 80 million × 2 paired-end reads (human, H1 ES cells) as a second dataset to evaluate our CRSP tool. total RNA was purified from RLT-Plus Buffer using RNeasy Plus Mini Kits (Qiagen, Netherlands) and quantified with both the Quant-It RNA Assay Kit (Thermofisher, Massachusetts) and the Bioanalyzer RNA 6000 Pico Kit (Agilent, California). The TruSeq RNA Library Prep Kit v2 (Illumina, San Diego), as per the manufacturer’s directions, was used to prepare and index all cDNA libraries. The RNA-seq library was sequenced on Illumina HiSeq 2500. We have submitted the paired-end RNA-seq data to GEO (accession number: GSE194237). We will make this dataset publicly available via NCBI GEO repository once the manuscript is accepted.

### 2.3 RNA-seq for Nile grass rat biopsies from 22 major organs

We collected 59 biopsy samples from 4 Nile grass rats representing 22 major organs (testis, brain, eye, pancreas, bone, thymus, bone marrow, spleen, heat, muscle, adipose, nerve, trachea, bladder, ovary, uterus, esophagus, stomach, liver, intestine, and kidney). Samples were prepared via the Illumina TruSeq RNA protocol. PolyA+ selection was undertaken via RNA purification beads followed by fragmentation for 6 minutes. We performed first and second strand cDNA synthesis and purified with Agencourt AMPure XP beads. We then performed an end repair step. We next adenylated the 3’ ends, ligated Illumina adapters and purified the libraries with AMPure XP beads. Then, we performed cDNA enrichment via PCR (15 cycles). Finally, we validated our libraries via PicoGreen quantification on a Synergy plate reader. Samples were then barcoded and sequenced as paired-end reads on the Illumina HiSeq 3000 sequencer.

### 2.4 Estimation of gene expression levels of Nile grass rat BodyMap and characterization of transcriptomic similarities of 22 major organs

The gene expression values of 59 Nile grass rat biopsy samples from 22 major organs were estimated by the CRSP tool. The transcriptomic contig set was mapped to the mouse protein database. The gene expression value was quantified as transcript per million (TPM) [2] (see **Supplementary Table S1:** TPM values for 59 biopsy samples), and then further normalized by media-of-ratio method implemented in EBSeq R package [13]. The gene expression value for each organ was calculated by the average of normalized TPMs in biological replicates. Pairwise Spearman’s rank-order correlations between these 22 organs were calculated to assess the transcriptomic similarity of these organs. The correlation-based dissimilarity matrix was used as the distance metric to perform a hierarchical clustering via average-linkage method. To assess the statistical uncertainty of the clustering, we computed two types of probability values: (a) bootstrap probability (BP) value which corresponds to the frequency that the cluster is identified in bootstrap copies, and (b) approximately unbiased (AU) probability values (p-values) by multiscale bootstrap resampling. A higher BP or AU value suggests more confidence that a cluster is less likely to be caused by random chance (e.g., clusters with AU and/or BP >0.95 are considered as high confident clusters). The hierarchical clustering and p-values calculation were performed via R package “pvclust” [14].

## 3. Results and Discussions

### 3.1 CRSP tool workflow and performance evaluations

As shown in **Figure 1A**, the basic workflow of CRSP includes merging assembly (contigs) from Trinity [5] tool, mapping contigs to a protein database, mapping RNA-seq reads to contigs, estimating contigs level expression values and imputing gene level expression values (see Methods for technical details).

**Figure 1:**
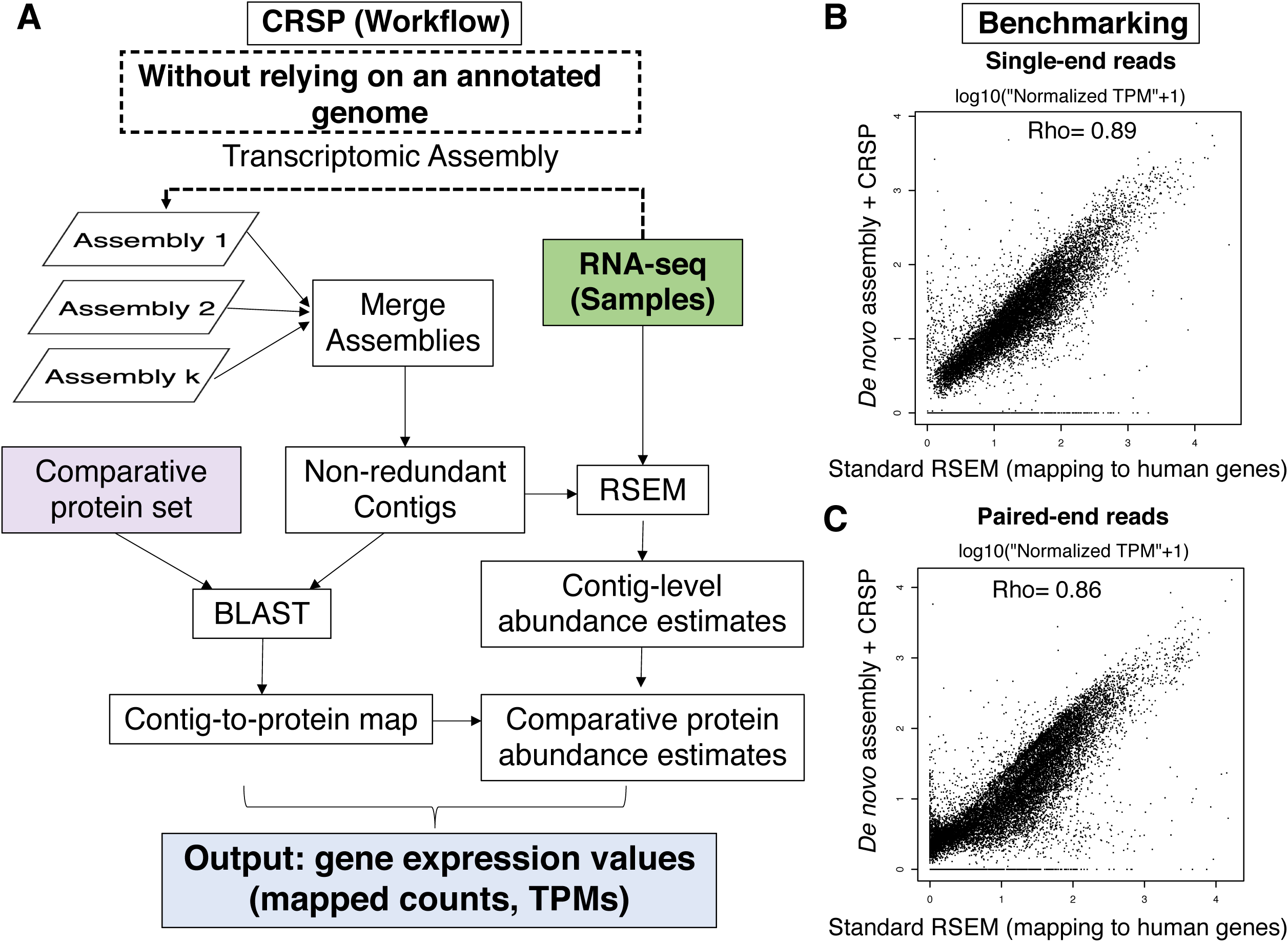
CRSP tool workflow and performance. (A) Workflow of CRSP; (B-C) CRSP estimated gene expression values are highly correlated with gene expression values estimated by directly mapping to a reference genome

It is widely accepted that gene expression values can be estimated by mapping deep read depth RNA-seq data to a well-annotated corresponding reference genome or gene sets. To generate a gold-standard gene expression value dataset to evaluate our CRSP tool, we pooled 578 million human H1 embryonic stem cells (ESCs) single-end RNA-seq reads from one of our prior studies [12]. We compared the gene expression values estimated by directly mapping to human reference genes via RSEM tool [2] and our CRSP tool (assuming there is no human reference genes or proteins available; using mouse proteins as reference instead). As shown in **Figure 1B**, the gene expression values which were estimated by CRSP are highly correlated with the gold-standard gene expression values (Spearman’s rank correlation coefficient Rho= 0.89). To further evaluate the performance of CRSP for paired-end RNA-seq data, we generated 80 million × 2 paired-end reads from human ESCs as a second dataset testing (see Methods). As shown in **Figure 1C**, the CRSP also achieved a high accuracy (Spearman’s Rho = 0.86) on this dataset.

### 3.2 Estimation of minimal RNA-seq read depth requirement to use CRSP

To further estimate the minimal reads requirement to run CRSP, we randomly sampled RNA-seq reads from pooled 578 million human ESCs single-end RNA-seq reads, and then estimated the Spearman’s Rho of gene expression values estimated from CRSP and from directly mapping to human reference genes, respectively. As shown in **Figure 2**, although the performance of CRSP progressively increased with more sequencing reads, 10-20 million single-end reads can be sufficiently to achieve reasonable accuracy. Importantly, if we have a pre-compiled transcriptomic contigs set from a deep sequencing (optional but not required), CRSP does not require deep read depth to achieve high accuracy (**Figure 2, blue line**). Together, our study suggests that CRSP is a useful tool making RNA-seq as a general tool to interrogate transcriptomic dynamics but not only limited to species with sequenced genomes or known transcripts.

**Figure 2:**
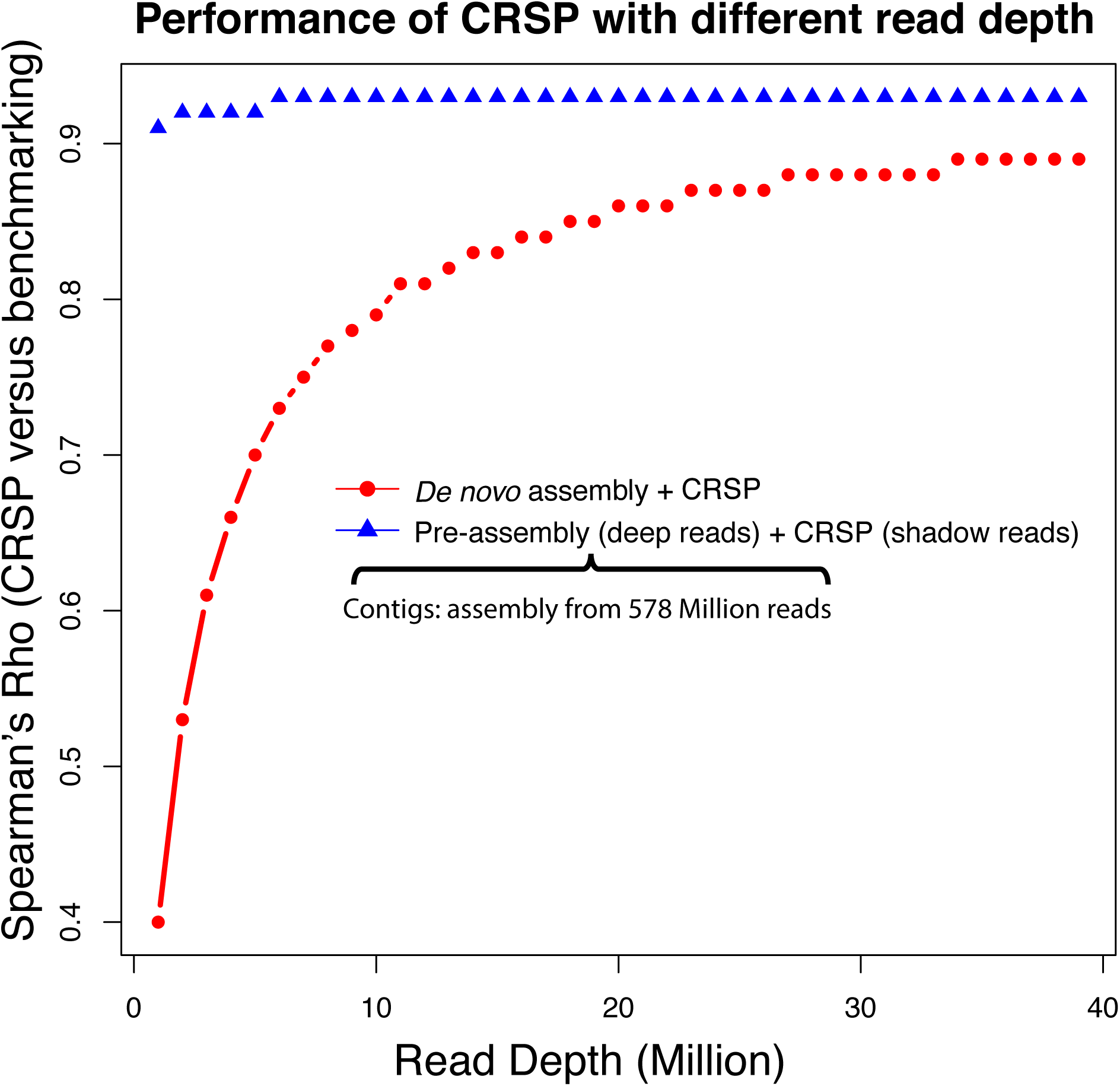
The performance of CRSP with different read depth.

### 3.3 Transcriptomic landscape of the Nile grass rat across 22 organs

Nile grass rat, as one of the most important type 2 diabetes models, has many unique characteristics. For example, instead of requiring high fat feeding, the higher caloric content relative to its natural diet (leaves and stems supplemented with insects) is sufficient to make these rats diabetic. The progression of diabetic retinopathy in the Nile grass rat [15], demonstrating its capacity to mimic a long-term diabetic complication with observable clinical features that were rarely reported in other type 2 diabetic rodent models. Hence, Nile grass rat diabetic model can accurately depict type 2 diabetes physiology in the human relative to other rodent models. However, there is very limited transcriptome studies on Nile grass rat. In this study, we generated an ultra-deep (2.3 billion × 2 paired-end) Nile grass rat RNA-seq data from 59 biopsy samples representing 22 major organs (testis, brain, eye, pancreas, bone, thymus, bone marrow, spleen, heat, muscle, adipose, nerve, trachea, bladder, ovary, uterus, esophagus, stomach, liver, intestine, and kidney). We used the CRSP tool to estimate the gene expression levels of these 59 biopsy samples and provide a unique resource for baseline spatial expression of Nile grass rat of major organs (**Supplementary Data**). To assess the organ similarity and differences, we applied hierarchical clustering based on all gene expression values (see Methods). As shown in **Figure 3**, the testis shows significantly distinguished overall gene expression patterns with other 21 organs.

**Figure 3:**
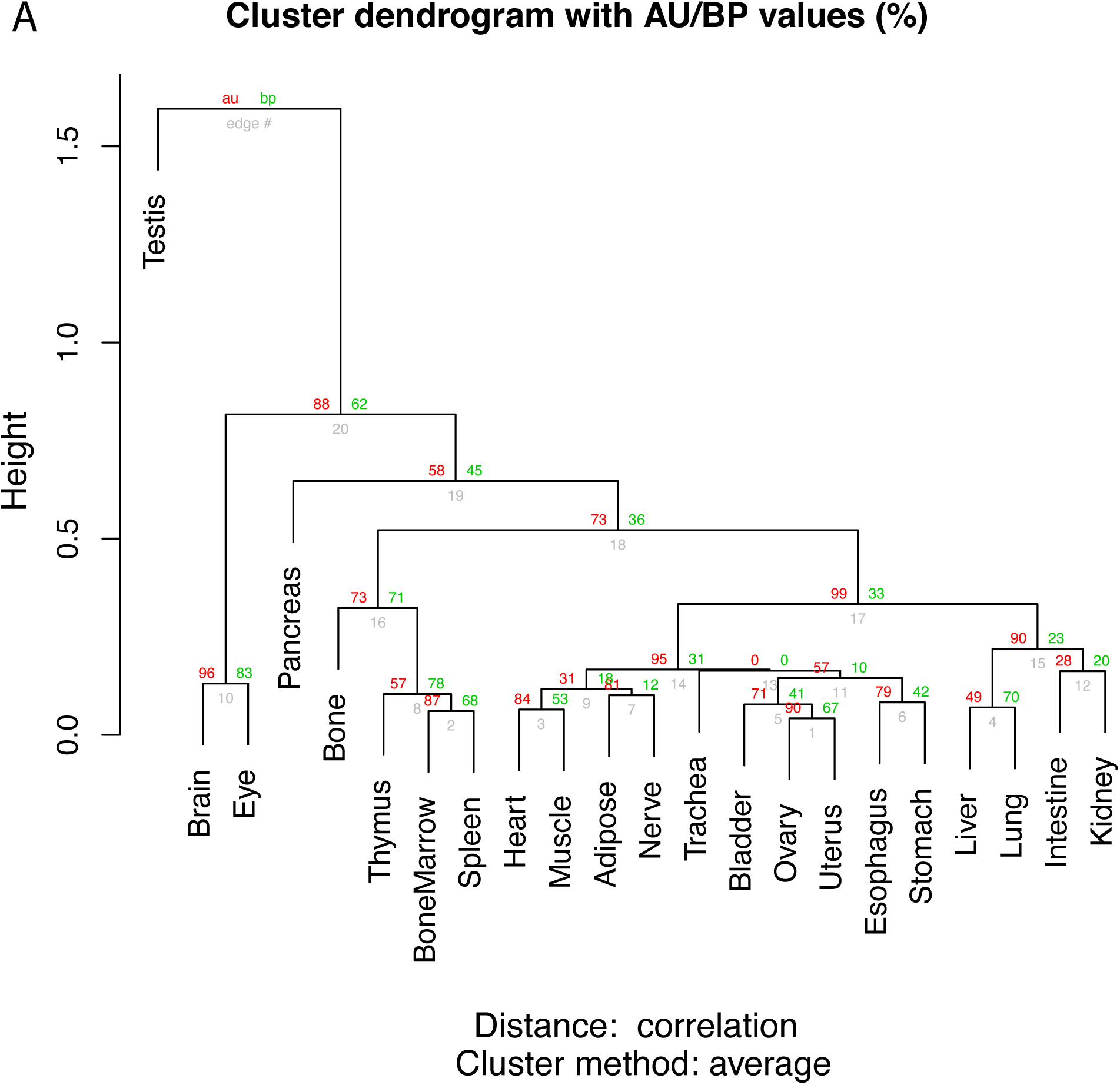
Transcriptomic similarities among 22 organs based on hierarchical clustering analysis.

Interestingly, our finding is consistent with a transcriptomic study on 16 human tissues (amnion, chorion, decidua, adipose, adrenal, brain, breast, colon, heart, kidney, liver, lung, lymph node, ovary, prostate, skeletal muscle, testes, thyroid and white blood cells) that testis had the greatest number of tissue specific genes[16], indicating that the gene expression patterns in testis are different from other tissue/organs in both human and Nile grass rat. In general, the tissue similarity in 22 major organs (**Figure 3**) is consistent with commonly accepted notion. For example, the muscle is transcriptomic similar to heart while brain and eye are similar but different with most of the rest organs.

### 3.4 Similarity and differences of baseline expressions in different central nervous system (CNS) regions of diurnal Nile grass rat

Nile grass rats have also been used to study the mechanisms underlying the impact of environmental lighting conditions on mood and cognitive functions as seen in humans [17]. The influence of light on mood is best exemplified by a condition termed as Seasonal Affective disorder (SAD), a major depressive disorder in which patients experience depression symptoms in fall and winter followed by spontaneous remission in spring and summer [18]. Bright light exposure has been shown to be effective in treating SAD and non-seasonal depression [18-21], suggesting light is a modulator of mood in humans. In addition, light is also a modulator for cognitive function, i.e. brighter illumination is associated with better cognitive performance and increase brain activity associated with the cognitive task [22, 23]. In Nile grass rats, it has been found that reducing daytime light intensity, i.e. a winter-like light condition, leads to increase depression-like behaviors including anhedonia and low libido as seen in depressed patients; as well as impairment in hippocampal-dependent spatial memory [17]. The results collectively underscore the validity of grass rats as a valuable model for elucidating the neural mechanisms underlying the modulatory role of light on mood and cognitive functions. A transcriptome wide analysis of genes and gene regulatory networks that are responsible for light-dependent changes in mood and cognition will contribute to fill a critical gap to understand how light conditions regulate our brain functions. As the first step toward this goal, we examined the expressions of a group of genes of interest, to analyzed the similarity and differences of their expression in a few brain regions and spinal cord of Nile grass rats. We selected fifteen genes that are known to be associated with light response or depressive behaviors based on our prior Nile grass rat studies [24, 25], genome-wide association study (GWAS) on Seasonal affective disorder (SAD) [26, 27], and transcriptome studies on various psychiatric and neurological disorders [28-34], to perform a hierarchical clustering analysis (**Figure 4**). The results have revealed that the caudal cortex and hippocampus (Hipp) are clustered while the pons, medulla and spinal cord are clustered with similar gene expression pattern. The relative expression patterns in different CNS regions are consistent with those in mouse or human brains reported on the Allen Brain Atlas, e.g., Bdnf and Zbtb20 are highest in the Hipp, Sst is lowest in the cerebellum. It suggests that although there are significantly differences between diurnal (e.g., human and Nile grass rat) and nocturnal (e.g., mouse) species regarding to temporal/circadian profile of gene expressions in the brain [35], the transcriptomic baseline similarity and differences in different brain regions remain largely conserved across diurnal and nocturnal species. The results support the feasibility of using the CRSP approach in Nile grass rats to address questions on molecular mechanisms underlying health and disease of central nervous system including psychiatric and neurological disorders.

**Figure 4:**
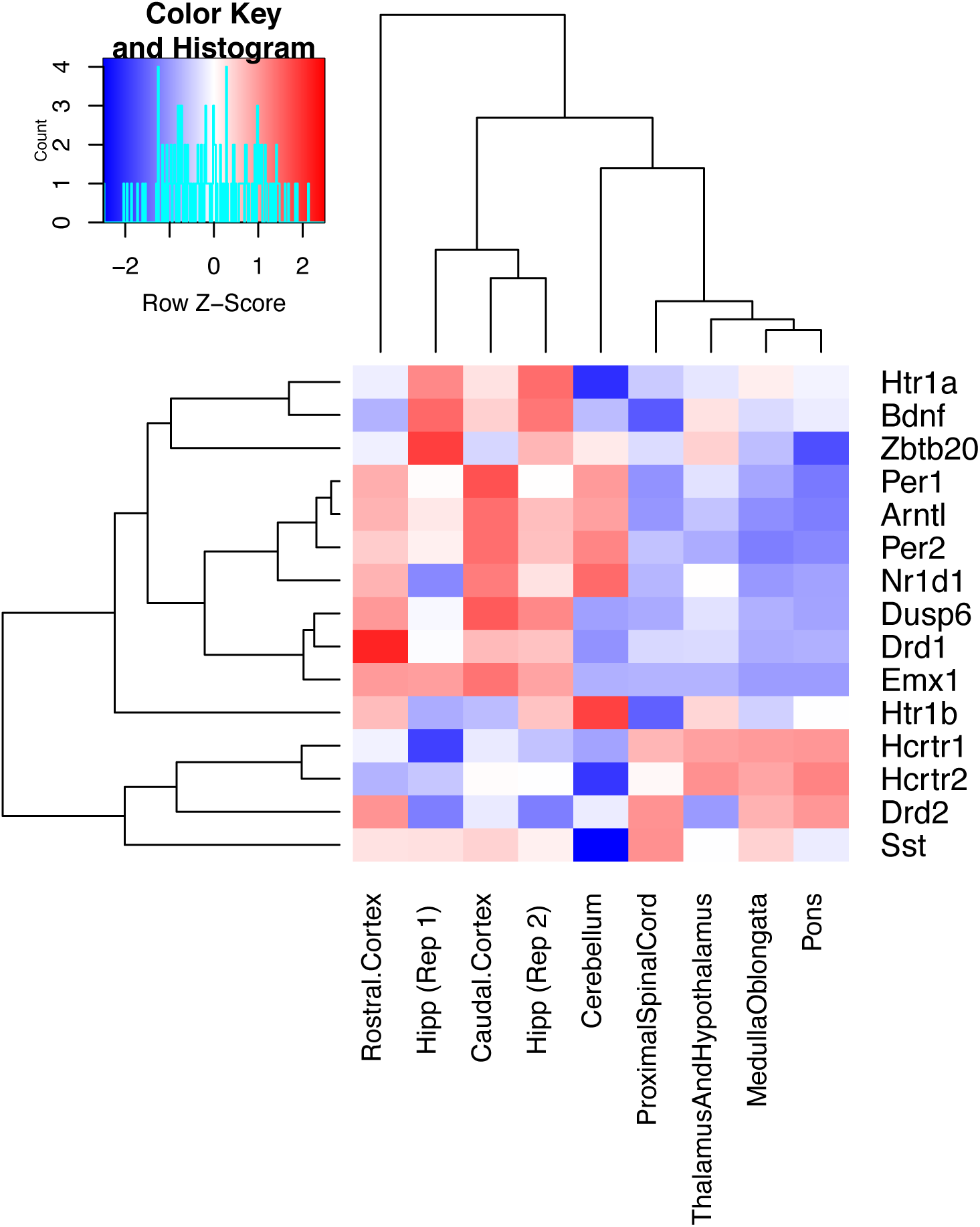
Similarity and differences of different regions in central nervous system (CNS) for expression of key light-response or depression marker genes in Nile grass rat.

## Conclusions

We developed CRSP, a comparative RNA-seq pipeline which allows estimation of gene expression values from RNA-seq data for species lacking a sequenced genome or corresponding reference transcripts. Benchmarking suggests that CRSP can achieve high accuracy for quantifying gene expression values. Applying CRSP to our generated ultra-deep (2.3 billion × 2 paired-end) Nile grass rat RNA-seq data from 22 major organs in this study, we provided a unique resource and spatial gene expression reference for using Nile grass rat as a model to study human diseases.

## Acknowledgment

This study was supported by the Garland Initiative for Vision funded by the William K. Bowes Jr. Foundation and Dr. Jiang’s startup funding in Cleveland State University.

